# Investigating *Cannabis sativa* L. gene expression through housekeeping genes and gene coexpression networks

**DOI:** 10.1101/2025.07.16.665086

**Authors:** Kevelin Barbosa-Xavier, Thiago M. Venancio

## Abstract

*Cannabis sativa* L. is a versatile crop with applications ranging from medicinal products to industrial materials. Despite growing interest in cannabis transcriptomics, comprehensive studies of gene expression in this species remain limited. Here, we explore cannabis transcriptomics through two complementary approaches: the identification and prioritization of housekeeping (HK) genes for qRT-PCR normalization and the construction of a genome-wide gene coexpression network (GCN) using publicly available RNA-Seq data. Based on expression stability criteria (i.e. log mean TPM ≥ 6, log variance < 0.3), we proposed the optimal HK genes and designed qRT-PCR primers for them. Primer specificity was verified *in silico* using the Jamaican Lion mother + Y genome. In parallel, to explore the non HK gene expression, we constructed a GCN comprising 32 coexpression modules, which was then analyzed for tissue-specific expression patterns, transcription factor (TF), and functional enrichment. Several modules were significantly correlated with cannabis tissues such as flowers and bast fibres and were linked to key processes, including photoperiod sensitivity, stress responses, and vegetative-to-reproductive phase transitions. Together, our results help understand the functions and maximize the use of HK genes in molecular biology and reveal coexpression modules that constitute critical genetic circuits underlying cannabis physiology.

## INTRODUCTION

Cannabis (*Cannabis sativa* L.) is a multifunctional crop that serves as the raw material for a variety of products, ranging from medicinal uses to construction materials and bioplastics (Gao et al. 2020). In recent decades, changing regulatory frameworks and a growing emphasis on sustainable agriculture have reignited interest in its cultivation and diversified applications. These changes have paved the way for advanced genomic studies, prompting the steady increase of cannabis RNA-Seq data in recent years, encompassing various plant tissues and biological processes, including secondary metabolite production (such as cannabinoids and terpenoids), fibre production, and plant defense mechanisms (Barbosa-Xavier et al. 2024).

Gene coexpression networks (GCNs) are widely used to investigate gene functions and interactions (Almeida-Silva et al. 2020). GCNs reveal groups of genes that exhibit similar expression patterns (i.e. modules) across conditions or plant tissues, enhancing our understanding of complex biological processes and gene interactions. On the other hand, genes that exhibit stable, high expression across all samples are likely to serve maintenance functions, making them housekeeping (HK) genes. The consistent expression of HK genes also make them good references or controls to normalize qRT-PCR results and facilitate comparisons of target gene expression (Gao et al. 2024, Parida et al. 2024).

Recently, our group processed and compiled publicly available cannabis RNA-Seq data to develop the Cannabis Expression Atlas (CannAtlas) (Barbosa-Xavier et al. 2024). Among other features, CannAtlas includes the identification of HK and tissue-specific genes, providing a valuable starting point for various expression analyses, including inferences from GCNs and the identification of reference genes for qRT-PCR analysis.

In this study, we explored the functions and designed primers for HK genes available in CannAtlas. We also constructed a GCN based on the expression of 27,640 genes across 394 samples available in CannAtlas and conducted functional enrichment analysis to find key transcription factors (TFs), Gene Ontology (GO) terms, and metabolic pathways overrepresented in coexpression modules. Finally, we also identified modules that are correlated with samples and discussed their relevance for cannabis biology.

## MATERIAL AND METHODS

### 1. Data collection

The normalized expression data in TPM (Transcripts Per Million) for 27,640 genes in 394 samples were downloaded from CannAtlas - https://cannatlas.venanciogroup.uenf.br/ (Barbosa-Xavier et al. 2024).

### 2. Identification of housekeeping qRT-PCR putative genes and primer design

We modeled the expression (in TPM) mean-variance relationship for all HK genes identified by Barbosa-Xavier et al. (2024). We selected genes with log expression variance lesser than 0.3 and log mean expression of at least 6 as putative reference genes for qRT-PCR analysis.

We used Primer3 v2.6.1 (Untergasser et al. 2012) to design putative qRT-PCR primers based on putative reference genes coding sequences (CDS) by using the parameters: PRIMER_OPT_SIZE = 20; PRIMER_MIN_SIZE = 18; PRIMER_MAX_SIZE = 25; PRIMER_OPT_TM = 60.0; PRIMER_MIN_TM = 57.0; PRIMER_MAX_TM = 63.0; PRIMER_MIN_GC = 30.0; PRIMER_MAX_GC = 70.0; PRIMER_MAX_POLY_X = 4; PRIMER_PRODUCT_SIZE_RANGE = 80 - 120; PRIMER_MAX_END_STABILITY = 9.0; PRIMER_MAX_SELF_ANY = 8.0; PRIMER_MAX_SELF_END = 3.0; PRIMER_PAIR_MAX_COMPL_ANY = 8.0. PRIMER_PAIR_MAX_COMPL_END = 3.0; PRIMER_NUM_RETURN = 5.

After primer design we perform a BLAST (Camacho et al. 2009) against the CannAtlas reference genome (Jamaican Lion Mother + Y) (Barbosa-Xavier et al. 2024, McKernan et al. 2020) to verify primer specificity using the parameters: -outfmt 6, -task blastn-short, - word_size 7, -perc_identity 100, -max_target_seqs 5, -max_hsps 5, -best_hit_overhang 0.1, - best_hit_score_edge 0.1, -dust no, -soft_masking false -qcov_hsp_perc 100.

We kept only the primer pairs with both forward and reverse sequences with perfect match on just one genome region by BLAST results.

### 3. Enrichment analysis

The enrichment analysis was conducted with Fisher exact test and p-values corrected with the Benjamini-Hochberg (BH) method in R. A threshold of 0.05 was applied to retain enriched pathways.

### 4. Coexpression analysis

#### a. Data processing

The R package BioNero (Almeida-Silva & Venancio 2022) was used to infer the cannabis GCN. To reduce noise, we excluded genes with mean TPM < 5. Further, samples were filtered using the standardized connectivity (Z.k) method by *ZKfiltering()* function using Pearson correlation. Samples with Z.k < ™2 were considered outliers and were filtered out. To further refine the data, we applied a principal component (PC)-based correction to account for potential confounding factors. This adjustment helps remove unwanted variation while preserving biologically relevant signals. We also performed a quantile normalization to ensure that expression values are comparable across different samples or conditions. As a result, gene expression values approximated a normal distribution, improving the reliability of downstream analyses (Almeida-Silva & Venancio 2022).

#### b. Exploratory analysis

The heatmap plot of sample correlations are obtained by the *plot_heatmap()* function in BioNero using Pearson correlation. Custom colors are set with the *custom_palette()* function of *plot_heatmap()*. A Principal Component Analysis (PCA) was performed with the function *plot_PCA()* (Almeida-Silva & Venancio 2022).

#### c. Gene co-expression network inference

BioNero uses the WGCNA algorithm (Langfelder & Horvath 2008) to infer GCNs. To identify the best β power that satisfies the network scale-free topology (sft) we use the *SFT_fit()* function with the arguments *net_type = “signed”* and *cor_method = “spearman”*. The network was inferred by the function *exp2gcn()* with the arguments *net_type = “signed”* and *cor_method = “spearman”* and the pre-computed *SFTpower =* 11. Hub genes were identified by using the function *get_hubs_gcn()*.

#### d. Gene coexpression network analysis

#### i. Module-trait associations

The function *module_trait_cor()* with the argument *cor_method = “spearman”* was used to calculate the correlation of modules and tissues (Almeida-Silva & Venancio 2022).

#### ii. Enrichment analysis

By using the sample metadata from CannAtlas (Barbosa-Xavier et al. 2024) and the function *module_enrichment()* from BioNero (Almeida-Silva & Venancio 2022), we inferred the module enrichment of Gene Descriptions (encoded proteins), Classification, Tissue, and TF. Enrichment analyses were conducted as described in the previous section.

## RESULTS AND DISCUSSION

### Comprehensive functional analysis and primer design of cannabis housekeeping genes for robust qRT-PCR normalization

HK genes are those with stable expression across all plant cells and that play key roles in maintaining cellular functions (Joshi et al. 2022). To explore the functions of cannabis HK genes proposed by Barbosa-Xavier et al. (2024), we performed GO and KEGG pathway enrichment analyses (Supplementary Tables Ia and Ib). The top 10 enriched GO terms for HK genes (Figure 1a) are primarily associated with proteasome activities (GO:0008540, GO:0043161, GO:0036402), transcription regulation (GO:0006359, GO:0003729, GO:0033962), kinase processes (GO:0005956, GO:0106310), phosphatase activity (GO:0019903), and the TCA cycle (GO:0006099), a result that was also mirrored by KEGG pathway enrichment (Figure 1b). These enrichment analyses emphasize the essential roles that cannabis housekeeping genes play in maintaining core cellular processes.

**Figure 1.**
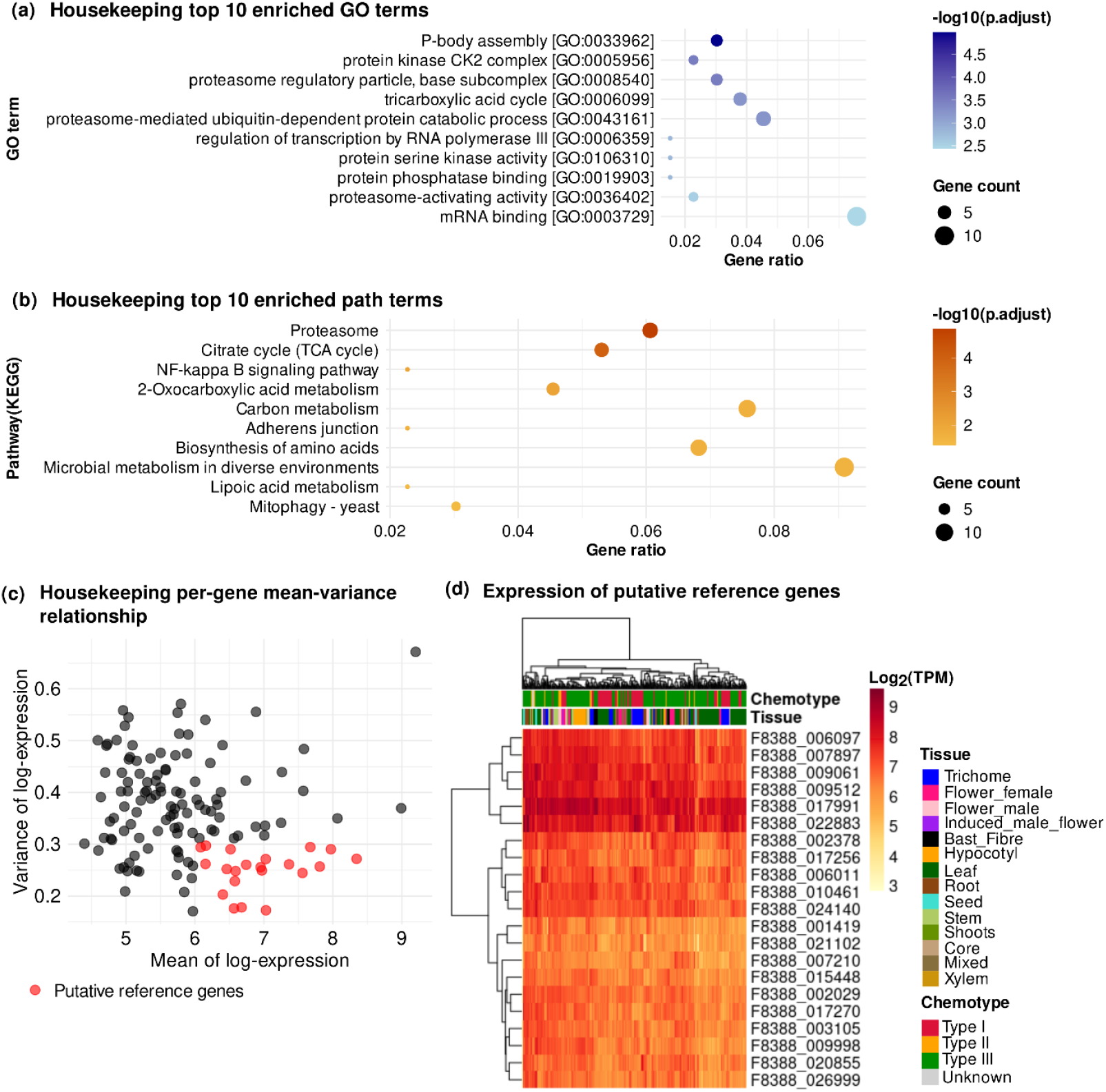
Comprehensive analysis of HK genes and proposed reference genes for qRT-PCR. (a) Gene ontology enrichment for HK genes, (b) metabolic pathway enrichment, (c) distribution of expression variance and selection of candidate reference genes, (d) expression patterns in CannAtlas samples.

Given the diversity of the data used in this study, which includes a wide range of samples, tissues, treatments, and varieties, we search for HK genes that would be suitable controls for qRT-PCR experiments. We selected the HK genes with log mean expression of at least 6 and variance lower than 0.3 (Figure 1c, 1d and Supplementary Table II) and designed primers for them (see methods for details). We design five primer pairs for each gene. To test primer specificity we perform a BLAST search of these primers to the Jamaican Lion mother + Y genome (as used on CannAtlas). By this test step we filtered no specific primer pairs, resulting in 67 putative qRT-PCR primer pairs for 17 HK genes (Supplementary Table III).

The average melting temperature (Tm) for the forward and reverse primers was 59.99°C (Supplementary Table III). The mean GC content for forward primers was 54.4%, and for reverse primers, it was 54.33% (Supplementary Table III). Primer Tm refers to the temperature at which half of the double-stranded DNA dissociates into single strands, while GC% indicates the proportion of guanine (G) and cytosine (C) nucleotides in the primer sequence. GC content between 40-60% and Tm temperatures between 50°C and 65°C are typically recommended for efficient annealing and optimal primer performance (Dieffenbach et al. 1993, Sharma 2020).

Interestingly, the candidate genes found here (Table I and Supplementary Table II) are not traditionally used for qRT-PCR normalization and are thus interesting novel alternatives. Five of these genes encode 20S and 26S proteasome subunits (Table I). Proteasomes are crucial for protein turnover in cells, functioning as a two‐part complex. The 20S core of the proteasome consists of stacked alpha subunits, which regulate substrate entry, and beta subunits, which contain proteolytic sites responsible for ubiquitinated protein degradation. The 26S proteasome regulatory particle includes AAA+ATPases that utilize ATP hydrolysis to recognize, unfold, and translocate substrates into the core for degradation. (Baumeister et al. 1998, Kurepa & Smalle 2008, Smith et al. 2006). Although proteasome activity can be influenced by stress conditions, this also is a basic cell maintenance function, which can justify the stable expression of some proteasome related genes. In addition, proteasome related genes were also found as potential qRT-PCR control genes by Chen et al. (2021) in strawberry.

**Table I.** Identification and descriptions of selected HK genes for qRT-PCR control.

| Gene_ID | Description | Definition(KEGG) |
| --- | --- | --- |
| F8388_001419 | Proteasome subunit alpha type | PSMA1; 20S proteasome subunit alpha 6 [EC:3.4.25.1] |
| F8388_009061 | AAA+ ATPase domain-containing protein | PSMC1, RPT2; 26S proteasome regulatory subunit T2 |
| F8388_009998 | PCI domain-containing protein | PSMD6, RPN7; 26S proteasome regulatory subunit N7 |
| **F8388_010461** | Proteasome subunit beta | PSMB3; 20S proteasome subunit beta 3 [EC:3.4.25.1] |
| **F8388_024140** | AAA+ ATPase domain-containing protein | PSMC6, RPT4; 26S proteasome regulatory subunit T4 |
| **F8388_006097** | V-type proton ATPase catalytic subunit A (EC 7.1.2.2) (V-ATPase 69 kDa subunit) (Vacuolar proton pump subunit alpha) | ATPeV1A, ATP6A; V-type H<sup>+</sup>-transporting ATPase subunit A [EC:7.1.2.2] |
| **F8388_007897** | Vacuolar proton pump subunit B (V-ATPase subunit B) (Vacuolar proton pump subunit B) | ATPeV1B, ATP6B; V-type H<sup>+</sup>-transporting ATPase subunit B |
| **F8388_017991** | V-type proton ATPase subunit E | ATPeV1E, ATP6E; V-type H<sup>+</sup>-transporting ATPase subunit E |
| **F8388_002029** | non-specific serine/threonine protein kinase (EC 2.7.11.1) | CSNK1, CKI; casein kinase 1 [EC:2.7.11.1] |
| **F8388_017256** | Casein kinase II subunit alpha (CK II alpha) (EC 2.7.11.1) | CSNK2A; casein kinase II subunit alpha [EC:2.7.11.1] |
| **F8388_017270** | Casein kinase II subunit alpha (CK II alpha) (EC 2.7.11.1) | CSNK2A; casein kinase II subunit alpha [EC:2.7.11.1] |
| **F8388_002378** | RRM domain-containing protein | TIA1, TIAL1; nucleolysin TIA-1/TIAR |
| **F8388_007210** | Ubiquitin domain; Heat shock chaperonin-binding | UBQLN, DSK2; ubiquilin |
| **F8388_021102** | Prefoldin subunit 5 | pfdA, PFDN5; prefoldin alpha subunit |
| **F8388_022883** | phosphoglycerate mutase (2,3-diphosphoglycerate-independent) (EC 5.4.2.12) | gpml; 2,3-bisphosphoglycerate-independent phosphoglycerate mutase [EC:5.4.2.12] |
| **F8388_026999** | CUE domain-containing protein |  |
| **F8388_015448** | Peroxisome biogenesis protein 22 |  |

In addition, three V-type proton ATPase subunits (Table I) were classified as qRT-PCR control candidates. V-type proton ATPases are vital for acidifying intracellular compartments, a function necessary for processes such as protein sorting, receptor-mediated endocytosis, and coupled transport (Holliday 2014). ATPases were also found as potential qRT-PCR control genes by Bharati et al. (2022) for *Salvia rosmarinus*.

Three Casein Kinase genes were also identified as potential controls for qRT-PCR analysis. Casein kinases are crucial for cell division by phosphorylating and destabilizing cyclin-dependent kinase inhibitors (Qu et al. 2021). Specific casein kinases also participate in light signaling pathways, flowering time regulation, and maintaining circadian rhythms (Lu et al. 2011, Qu et al. 2021). Studies in Arabidopsis thaliana show that genes encoding subunits of casein kinase II (CK2) are expressed ubiquitously across various tissues and developmental stages (Lu et al. 2011).

The last 6 genes proposed here (Table I) are the most intriguing. Although they fit our expression stability filter (see materials and methods) and have important functions in cellular maintenance, the literature indicates that the expression of these genes in other plant species are modulated by stressors (Cao 2016, Lorković & Barta 2002, Olmedilla & Sandalio 2019). The stability of these genes in cannabis may be a species-specific trait, but further studies are needed to confirm this hypothesis.

Although experimental proof is needed, the processes controlled by the HK genes proposed here are ubiquitously required and tightly regulated, which, together with their constant expression in various tissues, varieties and experimental conditions, reinforces their HK activity in cannabis and makes them particularly suitable for normalizing gene expression in qRT-PCR experiments.

### Unveiling gene coexpression networks and sample correlations in *Cannabis sativa*

To gain deeper insights into cannabis gene expression, we explored the correlations between samples and constructed a GCN. We used CannAtlas (Barbosa-Xavier et al. 2024) to retrieve TPM expression data for 27,640 genes and the metadata for 394 samples. After excluding genes with a mean TPM < 5 and filtering samples based on standardized connectivity (Z.k) < -2, 16,025 genes and 390 samples were kept for downstream analysis (Supplementary Table IV).

The resulting GCN consists of 32 modules (Figure 2a, b). The largest and smallest coexpression modules were coral2 (n = 1863) and yellow (n = 40), respectively, while the grey module, which contains unassigned genes, had 3,622 genes (Figure 2a). Pairwise correlations between module eigengenes revealed strong positive correlations in some modules (Figure 2c, Supplementary Table V). We also calculated the correlations between modules and tissues (Figure 3, Supplementary Table VI) and observed that each module was both positively and negatively correlated with multiple tissues.

**Figure 2.**
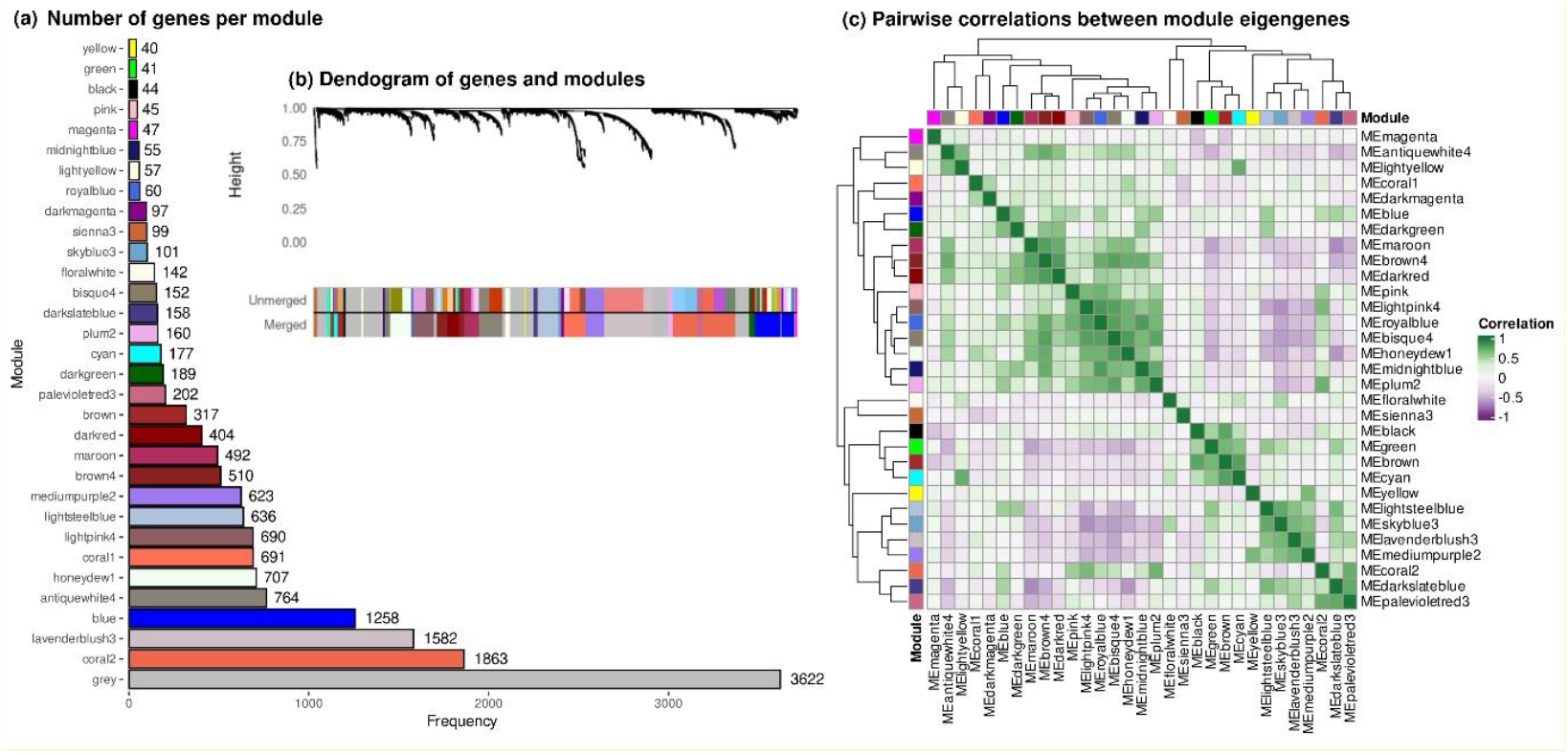
Cannabis gene co-expression network **(a)** number of genes per module, **(b)** dendrogram of genes and modules and **(c)** pairwise correlations between module eigengenes.

**Figure 3.**
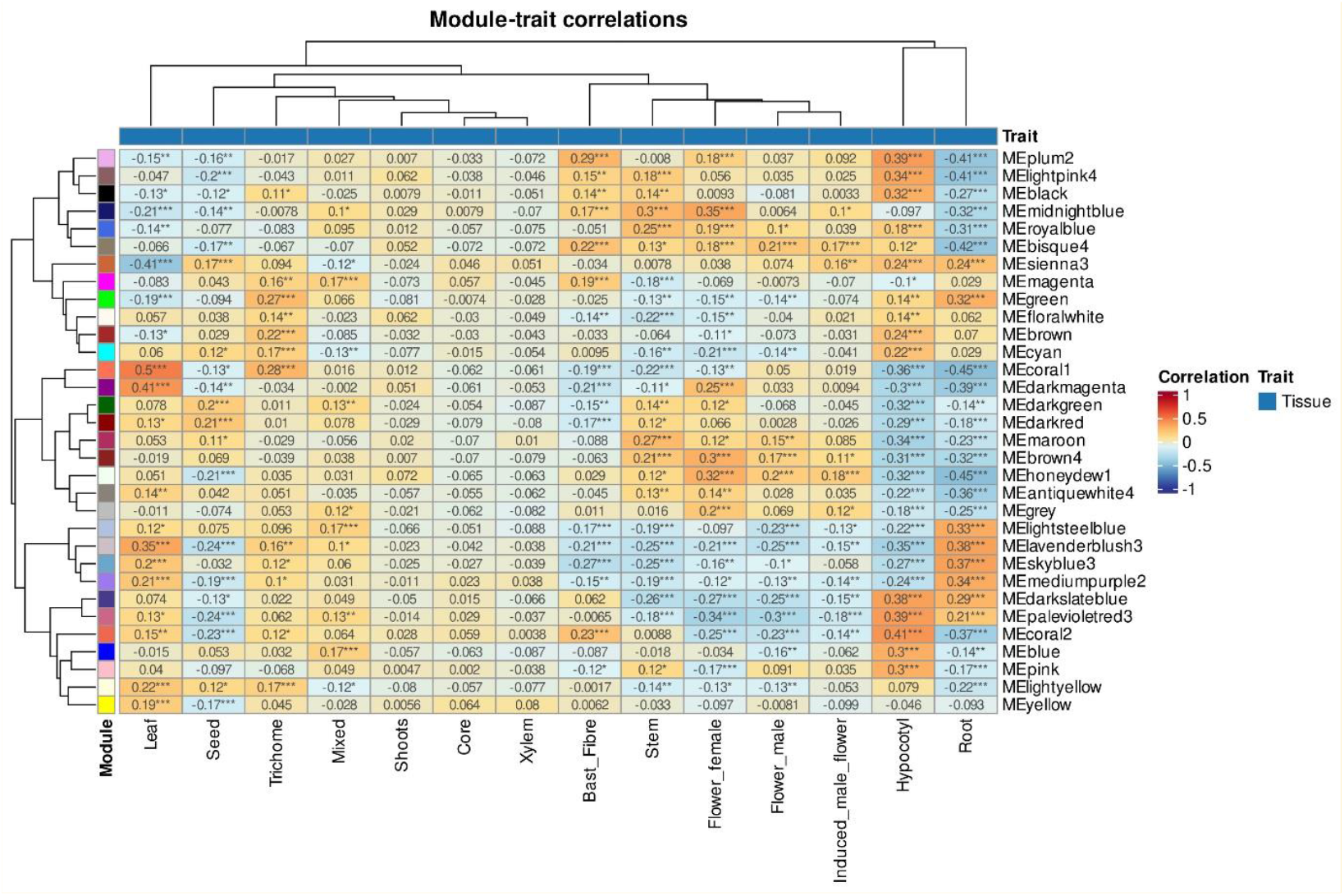
Correlation between coexpression modules and tissues.

To gain a deeper understanding of the overall functions of the modules, we performed enrichment analysis based on gene annotations provided by Barbosa-Xavier et al. (2024). The analysis focused on gene description, expression class, tissue-specificity, TFs, GO (UniProt), KEGG metabolic pathways, and InterPro domains (Supplementary Tables VII–XIII).

Pairwise sample correlations based on gene expression revealed interesting positive correlations in clades 1, 2, and 4 (Figure 4). Clade 1 groups samples from male (14 out of 18), female (22 out of 34), and induced male flowers (6 out of 6), alongside stem (17 out of 26), bast fibres (4 out of 12), and leaf (11 out of 151) samples. Clade 2 includes root (29 out of 32), mixed (8 out of 19), and leaf (36 out of 151) samples. Clade 4 brings together bast fibres (6 out of 12), hypocotyl (24 out of 24), stem (3 out of 26), leaf (1 out of 151), and core (1 out of 1) samples.

**Figure 4.**
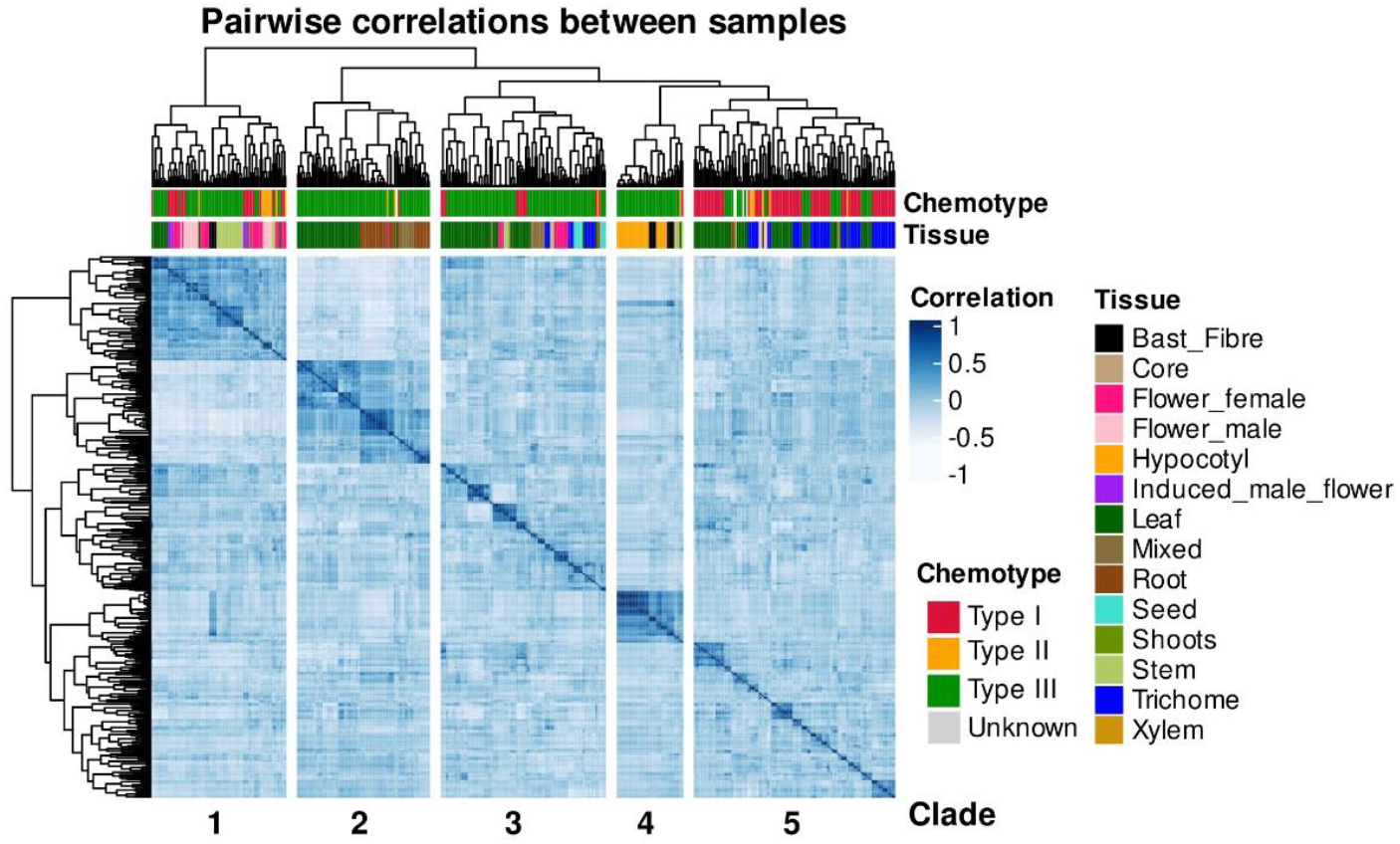
Cannabis samples pairwise correlation based on expression.

#### a. Clade 1 tissues are positively correlated with coexpression modules involved in vegetative to reproductive phase transition

The majority of stem (n = 15) and leaf (n = 8) samples in clade 1 (Figure 4) originate from a study of photoperiod-insensitive flowering and were collected across vegetative, pre-flowering, and flowering stages (Dowling et al., 2024; Bioproject PRJNA956491). The module-tissue correlation analysis indicates that the modules most positively associated with the primary tissues in clade 1 (flowers and stem) include midnightblue, royalblue, bisque4, sienna3, maroon, brown4, honeydew1, and antiquewhite4 (Figure 3). Additionally, pairwise module correlations (Figure 2c) reveal a strong correlation (>0.6) between midnightblue, royalblue, bisque4, and honeydew1 modules (Supplementary Table V). These modules contain 941 genes with at least 0.5 of expression correlation (Figure S1, Supplementary Table XV), including 46 genes classified as group-enriched or tissue-specific for tissues within clade 1 (Figure 4, Supplementary Table XIV).

The honeydew1 module is the biggest one on this GCN and is the main responsible for the functional enrichment observed. It is enriched for TF families SBP (Squamosa-Promoter Binding Protein), YABBY, and ZF-HD (Zinc Finger-Homeodomain). Additionally, it is associated with several protein descriptions, including the Chloramphenicol Acetyltransferase-like domain superfamily, DUF4005 domain-containing proteins, GDSL esterase/lipases, histones (H2A, H2B, H3, H4), SBP-type domain-containing proteins, and ZF-HD dimerization-type domain-containing proteins (Supplementary Tables VIII and X).

The gene set in clade 1 includes eight SBP TFs, which belong to a plant-specific family that regulate the MADS-box SQUAMOSA and play roles in early flower development (Klein et al. 1996). SBP genes have also been linked to the transition from vegetative to reproductive phases, hormone signal transduction, and leaf development (Gandikota et al. 2007, Jung et al. 2011, Usami et al. 2009, Zhang et al. 2007). Further, the SBP gene SPL8 regulates gibberellic acid (GA) biosynthesis and signaling genes, acting as a regulator of GA-dependent developmental processes in *Arabidopsis* (Zhang et al. 2007).

Additionally, these modules contain five ZF-HD TFs, which are known to play a role in biotic and abiotic stress responses, including salinity, drought, and temperature fluctuations. They are also involved in the regulation of flower development by mediating signaling responses to flowering hormones such as GA and 6-benzylaminopurine (6-BA) (Shalmani et al. 2019).

YABBY TF genes were also enriched in these modules. The YABBY TF family is plant-specific and regulates flowering and vegetative development processes, such as the formation of vascular tissues, lateral organs, carpels, male florets, leaves, ovule exoderm, and epicarp (Strable et al. 2017, Strable & Vollbrecht 2019, Villanueva et al. 1999, Yang et al. 2016). YABBY TFs also regulate GA biosynthesis and metabolism and contribute to abiotic stress responses, particularly in salinity and drought tolerance (Li et al. 2019, Zhao et al. 2017).

The GDSL gene family plays a crucial role in lipid metabolism, cell wall modifications, and secondary metabolism. It has been implicated in various developmental processes, including seed germination, coleoptile elongation, plant height regulation, fertilization, and the development of roots, stomata, floral organs, anthers, pollen, fruit, and seeds in several plant species, such as rice, maize, *Arabidopsis*, and soybean (An et al. 2019, Ishiguro et al. 2001, Shen et al. 2022, Su et al. 2020, Turquetti-Moraes et al. 2025, Yu et al. 2020). Additionally, chloramphenicol acetyltransferase-like domains serve as structural components of shikimate O-hydroxycinnamoyltransferase (HCT) proteins (D’Auria 2006). HCT plays a key role in lignin metabolism, supporting plant growth and enhancing cell wall resistance to biotic and abiotic stresses. Studies have shown that HCT expression responds to photoperiod changes, being down regulated under short-day conditions (Besseau et al. 2007, Ren et al. 2020). This reduction in lignin biosynthesis halts vegetative growth while promoting flavonoid accumulation and flowering (Besseau et al. 2007, Ren et al. 2020). Furthermore, histones H2A, H2B, H3, and H4, along with their variants, are essential for epigenetic regulation, influencing transcriptional response to environmental and biochemical cues (Xu et al. 2018). For example, the histone variant H2A.Z has been shown to regulate flowering by controlling the expression of FLOWERING LOCUS C (FLC) genes (Xu et al. 2018). It also plays a role in plant growth and development by modulating the expression of specific microRNAs, such as miR156A and miR156C (Xu et al. 2018).

Cannabis is classified as a short-day plant, meaning its reproductive phase is triggered when the photoperiod ranges from 11 to 15 hours of light, except for certain photoperiod-insensitive strains, commonly known as auto-flowering varieties (Schilling et al. 2023). The enrichment analysis of modules associated with clade 1 samples suggests that flower and stem tissues share key pathways and mechanisms related to the transition from the vegetative to reproductive phase, stress defense, and hormone signaling. Additionally, the enrichment of histones highlights a significant role for transcriptional epigenetic regulation within these modules. Notably, the low expression of miR156A and miR156C, both regulated by a H2A histone variant, has been linked to the vegetative-to-reproductive phase transition in *Arabidopsis* through the promotion of SBP gene expression (Xu et al. 2018). This further reinforces the association between the genes in these modules and phase transition processes in *C. sativa*.

#### b. Clade 2 tissues reveal coexpression modules associated with stress response and photoperiod-related pathways

The leaf samples in clade two (Figure 4) also originate from the Dowling et al. (2024) study on photoperiod response. Among the 29 root samples in this clade, 24 are from NaHCO_3_ stress studies (Cao et al., 2021, 2023; BioProjects PRJNA672722 and PRJNA813212). Additionally, the eight mixed samples belong to a cadmium stress study (BioProject PRJNA701120; Yin et al., 2022). The modules showing a positive correlation with leaf, root, and mixed samples in this clade include lightsteelblue (n = 636), lavenderblush3 (n = 1582), skyblue3 (n = 101), mediumpurple2 (n = 623), darkslateblue (n = 158), and palevioletred3 (n = 202) (Figure 3). Pairwise correlation analysis between modules indicates strong correlations among lightsteelblue, lavenderblush3, skyblue3, and mediumpurple2, while darkslateblue and palevioletred3 are closely correlated with each other but show lower correlation with the other modules (Figure 2c, Supplementary Table V).

These modules contain 3,223 genes with at least 0.5 of expression correlation (Figure S2, Supplementary Table XVI). Among these, 706 genes are either group-enriched or tissue-specific, with 512 being root-specific (Supplementary Table XVII). All root-specific genes belong to lavenderblush3, while root group-enriched genes are distributed across the lavenderblush3, mediumpurple2, and palevioletred3 modules (Supplementary Table XVII).

Given that the clade 2 samples originate from stress studies, the enrichment of these modules in plant defense-related pathways is expected. Our KEGG pathway enrichment analysis aligns with the findings of Cao et al. (2021 and 2023), which indicate that genes involved in hemp response to NaHCO_3_ stress primarily requires the following pathways: Phenylpropanoid biosynthesis, Plant hormone signal transduction, Valine, leucine and isoleucine degradation, Nitrogen metabolism, Plant-pathogen interaction, alpha-Linolenic acid metabolism, Arginine and proline metabolism (Supplementary Table XIII).

Cao et al. (2023) observed that the miR156 microRNA family is upregulated in hemp salt-alkali tolerant plants, enhancing the plant adaptability to stressful environments. This is particularly interesting because, as mentioned earlier, miR156 is also involved in epigenetic regulation in response to photoperiod changes, where it regulates genes like SBP that coordinate the transition from vegetative to reproductive phases through flowering induction (Xu et al., 2018). Based on this, the observed correlation between root and leaf samples suggests that both stress and photoperiod responses may control overlapping pathways that prevent flowering until environmental conditions are favorable.

#### c. Coexpression module 4 regulate transcription in hemp bast fibre and hypocotyl development

The hypocotyl and bast fibre samples belong to the bioprojects PRJNA436945 and PRJNA435671, and are derived from studies focused on hemp fibre development (Behr et al. 2016, Guerriero et al. 2017). According to Behr et al. (2016), hemp hypocotyl cells undergo primary and secondary growth phases between 6 and 20 days after sowing (DAS). From 6 to 9 DAS, cells are in the primary growth phase, marked by cell elongation, while from 15 to 20 DAS, they enter the secondary growth phase, characterized by secondary cell wall deposition.

The modules most strongly correlated with the tissues from clade 4 (hypocotyl and bast fibre) (Figure 4) are plum2, lightpink4, black, bisque4, and coral2 (Figure 3). Among these, lightpink4, bisque4, and plum2 modules show a stronger correlation with each other than with coral2. The black module, on the other hand, exhibits the lowest correlation with all the other modules (Figure 2c, Supplementary Table V). These modules contain 2.801 genes with at least 0.5 of expression correlation (Figure S3, Supplementary Table XVIII). Among these, 354 genes are group-enriched or tissue-specific, including 233 hypocotyl-specific genes, 22 bast fibre-specific genes, and 56 bast fibre/hypocotyl group-enriched genes (Supplementary Table XIX). Nearly all of the hypocotyl/bast fibre tissue-specific or group-enriched genes are found in the coral2 module (n = 305), while the remainder are distributed across the lightpink4 (n = 5) and plum2 (n = 1) modules (Supplementary Table XIX).

The network contains 15 hub genes that are TFs, with 14 from the coral2 module and one from the plum2 module (Figure S3). Among these, eight TFs are hypocotyl-specific (n = 7) and bast fibre-specific (n = 1), all from the coral2 module (Supplementary Table VII). The TFs include: four NAC (F8388_003104, F8388_003739, F8388_017539, F8388_026233), three MYB (F8388_005875, F8388_009800, F8388_016389), two TALE (F8388_012375, F8388_014869), two MYB-related (F8388_021332, F8388_023517), one WOX (F8388_001731), one HD-ZIP (F8388_001015), one HB-other (F8388_016512), and one G2-like (F8388_025213) (Supplementary Table VIII). Genes from all these TF families have been linked to bast fibre development in the studies by Behr et al. (2016) and Guerriero et al. (2017).

Our GCN analysis unraveled a high coexpression of genes involved in secondary cell wall deposition in hypocotyl, inducing the bast fibre development, supporting the results from Behr et al. (2016) and Guerriero et al. (2017) findings. However, studies involving gene expression at different hemp varieties focusing not just into fibre development but also into fibre quality are yet to be conducted, in particular if we consider the range of industrial applications of cannabis bast fibres.

## CONCLUSIONS

Although lab tests need to be performed to confirm the suitability of the proposed genes to qRT-PCR control, the use of a diversified expression dataset as the Cannabis Expression Atlas (Barbosa-Xavier et al. 2024) provides a promising identification of robust HK genes.

Through coexpression analysis, we identified key gene modules associated with cannabis vegetative to reproductive phase transition, photoperiod response, and stress response pathways. We found that flower and stem samples were closely linked to modules regulating pathways involved in phase transition and hormonal signaling, while root and leaf samples under stress uncovered modules related to plant defense and environmental adaptation, including photoperiodicity. The overlap between photoperiod sensitivity and stress response pathways suggests shared regulatory mechanisms that control flowering under suboptimal conditions. Additionally, our analysis of hypocotyl and bast fibre samples in clade 4 revealed gene coexpression modules driving primary and secondary growth. Modules like coral2, enriched for fibre-specific genes and TFs such as NAC, MYB, and WOX, are crucial for secondary cell wall deposition and bast fibre development. Overall, our gene coexpression network analysis provides a valuable resource for future functional studies on *C. sativa* development and stress response. This knowledge can inform breeding strategies aimed at optimizing traits such as fibre quality, flowering control, and stress tolerance, ultimately enhancing the economic and agronomic potential of hemp varieties.

## ACKNOWLEDGMENTS

This work was supported by Fundação Carlos Chagas Filho de Amparo à Pesquisa do Estado do Rio de Janeiro (FAPERJ), Coordenação de Aperfeiçoamento de Pessoal de Nível Superior-Brasil (CAPES; Finance Code 001), and Conselho Nacional de Desenvolvimento Científico e Tecnológico (CNPq). The funding agencies had no role in the design of the study and collection, analysis and interpretation of data and in writing.

## AUTHOR CONTRIBUTIONS

Conceived the study: K.B.-X. and T.M.V.; Funding and resources: T.M.V.; Data analysis: K.B.-X.; Interpretation of the results: K.B.-X. and T.M.V.; Wrote the manuscript: K.B.-X. and T.M.V.

