## Supplementary figures for "Investigating *Cannabis sativa* L. gene expression through housekeeping genes and gene coexpression networks"

Kevelin Barbosa-Xavier - https://orcid.org/0000-0002-3750-5331
Thiago Motta Venancio - https://orcid.org/0000-0002-2215-8082

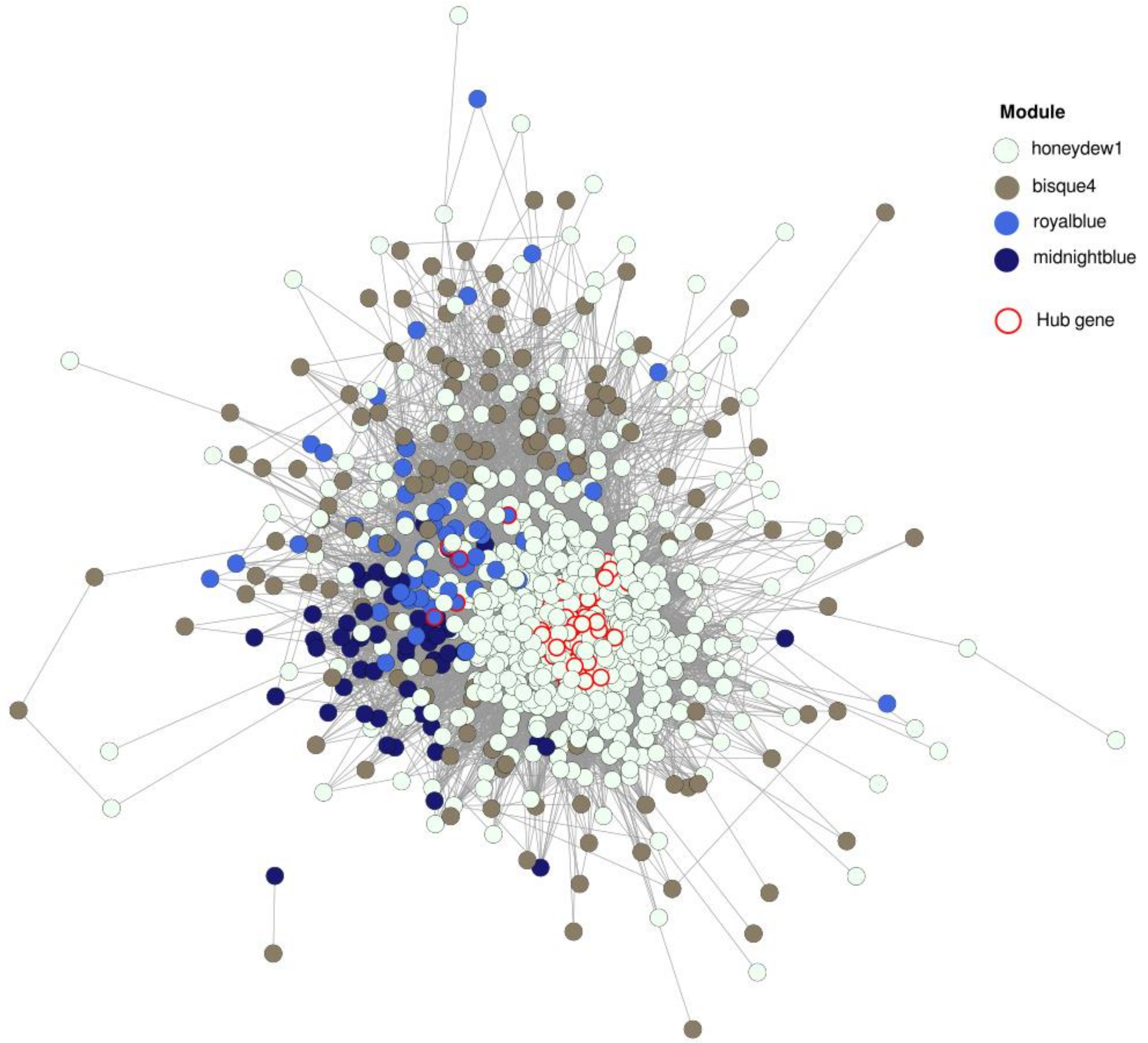


**Figure S1.** Coexpression network of Clade 1 genes. Nodes represent individual genes, and edges indicate coexpression relationships weighted by interaction strength. Genes are colored according to their module assignment. Red-bordered nodes denote hub genes.

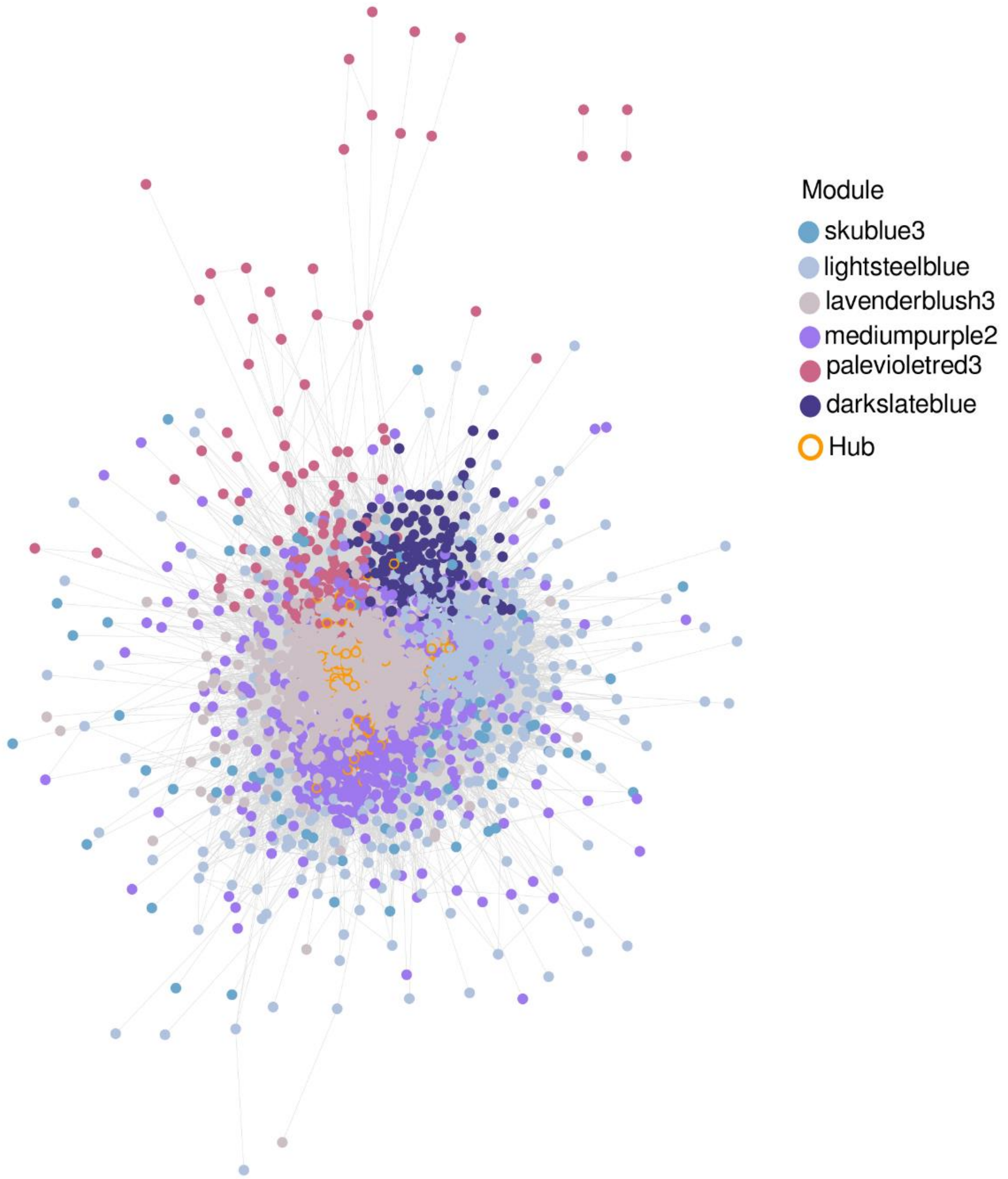


**Figure S2.** Coexpression network of Clade 2 genes. Nodes represent genes, and edges represent coexpression relationships weighted by interaction strength. Node colors indicate module membership. Hub genes are highlighted with orange borders.

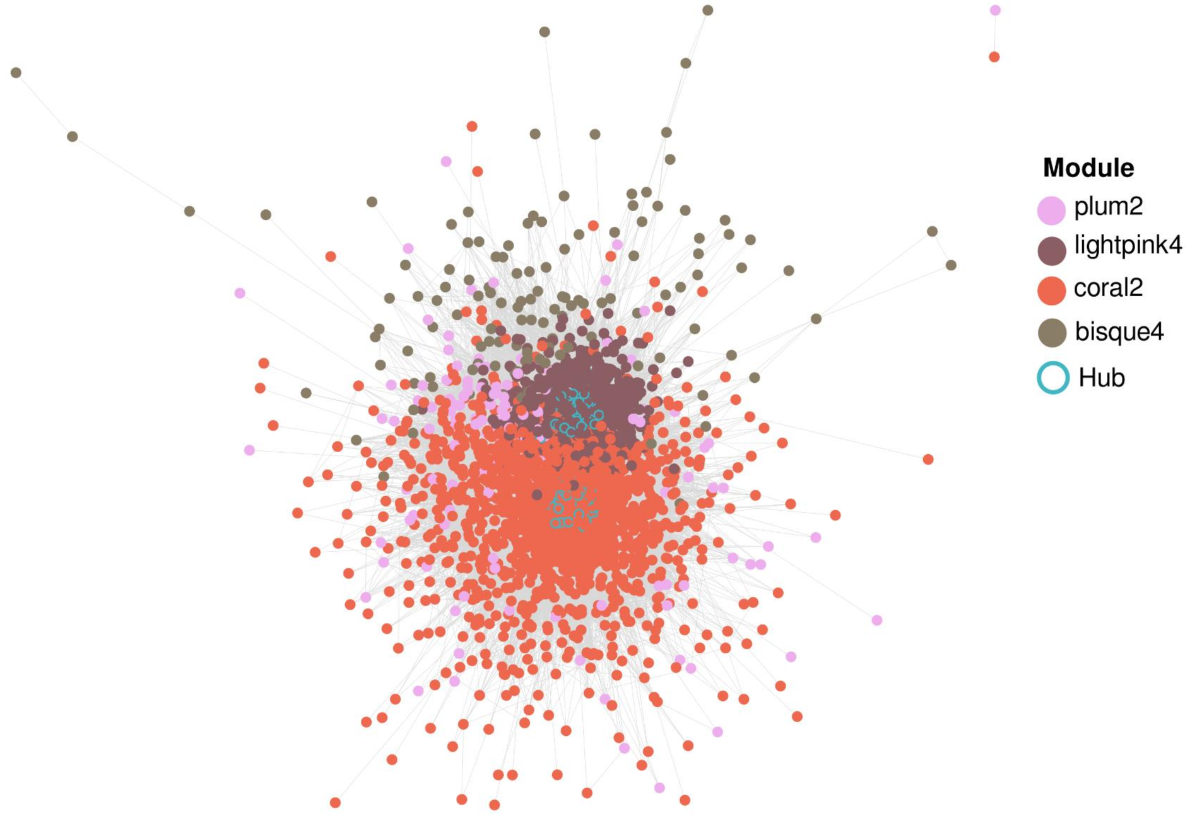


**Figure S3.** Coexpression network of Clade 4 genes. Nodes represent genes, and edges represent coexpression relationships weighted by interaction strength. Node colors indicate module assignment. Hub genes are highlighted with blue borders.
